# Single-cell RNA sequencing of peripheral blood cells identifies transcriptomic signatures in Parkinson’s disease

**DOI:** 10.64898/2026.03.30.715379

**Authors:** KC Paul, O Wilkins, E Carloni, E Fikse, LA Salas, S Lee, M Feldman, R Thompson, GE Kersey, CA Jeffreys, F Kolling, DM Kasper, JK Lee, M Havrda

**Affiliations:** Department of Molecular and Systems Biology, Geisel School of Medicine at Dartmouth, Hanover, NH; Dartmouth Cancer Center, Dartmouth-Hitchcock Medical Center, Lebanon, NH; Department of Epidemiology, Geisel School of Medicine at Dartmouth, Hanover, NH; Department of Neurology, Dartmouth-Hitchcock Medical Center, Lebanon, NH; Department of Physiology and Pharmacology, College of Veterinary Medicine, University of Georgia, Athens, GA; Department of Biomedical Sciences, University of New England College of Osteopathic Medicine, Portland, ME

**Keywords:** Parkinson’s disease, peripheral immune cells, scRNA-Seq, transcriptomics, immune dysregulation

## Abstract

Parkinson’s disease (PD) is a progressive age-related neurodegenerative disorder characterized by both motor and non-motor symptoms. The poorly understood prodromal period, decades-long progression, and disease-phenotype heterogeneity continue to impede the development of preventive and curative therapies. A growing appreciation of immune system changes during the progression of PD suggests that evaluating peripheral immune cells may help identify signatures relevant to disease etiology. We employed single-cell RNA sequencing to profile the transcriptomes of peripheral blood mononuclear cells (PBMCs) from a cohort of 12 patients with PD and 12 healthy controls, equally distributed by sex. Analysis identified gene expression signatures specific to immune cell lineages in PD when compared with healthy controls. Analysis of the dataset indicated that aspects of the PD-related changes were associated with sex, including metabolic and inflammatory changes. Further analysis of myeloid and T cell subsets identified additional pathways and gene expression profiles associated with PD. Trajectory analysis of the myeloid and T cell datasets indicated significant changes in the distribution of cells across states of gene expression in PD compared with controls. This work provides new evidence of peripheral immune cell changes in PD utilizing high-resolution transcriptomics in a cohort powered to analyze sex as a variable.

**Highlights:** Transcriptomic dataset in a cohort powered to analyze immune phenotype in Parkinson’s disease

Parkinson’s disease-specific gene expression signatures in peripheral immune cell lineages Identification of sex differences in the immune cell transcriptome in Parkinson’s disease

Trajectory analysis identifies changes in immune cell phenotypic distribution in Parkinson’s monocytes

## Introduction

Parkinson’s disease (PD) is a molecularly complex pathology that progresses over decades and in which degeneration of neurons in the central nervous system (CNS) results in debilitating motor deficits. Molecular and cellular changes detected throughout the slow progression of PD suggest that early detection and neuroprotection may be possible. Unfortunately, despite a growing understanding of PD biology, only symptomatic treatments are available. There is no single cause for PD, but rather complex gene-environment interactions contribute to risk, incidence, and progression^1,2^. Genetic mutations identified in familial PD highlight the importance of protein misfolding, mitochondrial function, redox homeostasis, and vesicular transport^3^. Large-scale genome-wide association studies continue to identify new risk loci, many of which are associated with the immune response, and mendelian randomization analyses have identified immune mediators in PD pathogenesis^4,5^.

The peripheral immune system interfaces with the brain and can sense and respond to neurological damage and distress^6,7^. Under normal circumstances, peripheral immune cells patrol and survey the vascular endothelium and can enter tissues, differentiate, and perform a variety of homeostatic and repair functions^8–10^. Multiple genetic and immunological studies indicate that cells of the peripheral mononuclear phagocyte system are consistently altered in PD (for review ^11–14^), especially monocytes, which enter the brain and contribute to neuroinflammation^15–18^. A reduced proportion of extravasation-competent CD16+ monocytes compared with CD14+ monocytes is observed in PD^19,20^ and other neurodegenerative diseases^14^. In mouse models, border-associated macrophages (BAM) residing in the neuronal perivascular space orchestrate the recruitment of peripheral immune cells into the brain parenchyma, driving alpha-synuclein (α-syn) induced neurodegeneration^21^. The same study noted that in patients, BAM-rich regions of the PD brain were enriched for T cells. In concordance, a longitudinal case study identified elevated α-syn-specific T cell responses detected prior to the diagnosis of motor PD^22^. These studies support continued efforts to increase the understanding of the peripheral immune contribution to PD based on their potential to alter CNS homeostasis, augment neurotoxic inflammation, and impact the progression of PD^15,18,23–25^.

Genome-wide association studies (GWAS) in large cohorts of PD patients and controls support transcriptomic screening of immune cells by identifying polymorphisms in genes associated with the immune response. Meta-analysis of datasets containing PD patients and healthy controls identified risk loci with probable impact on genes directly associated with immune function, including *NFKB2* and *NOD2*, important mediators of the host response to pathogens and damage-associated molecular patterns (DAMPs)^26^. The same study also identified numerous other loci predicted to impact genes expressed in immune cells, particularly myeloid lineages, including *SCARB2*, *VAMP4*, and *LRRK2*. Recent studies further support the use of high-resolution transcriptomics to identify PD-related changes in peripheral immune cells. Moquin-Beaudry et al. identified differentially expressed genes across immune lineages in a PD cohort and observed gene expression profiles indicating a phenotypic shift towards an activated state in monocytes and altered proportions of lymphocyte subtypes utilizing single-cell RNA sequencing (scRNA-Seq)^27^. Xiong and others utilize scRNA-Seq to identify gene expression changes associated with PD severity and distinct NK cell phenotypes in PD compared with controls^28^. There is also previous work from others that describes sex-related immune cell phenotypic differences in PD and other neurodegenerative disorders^29,30^. Carlisle et al. identified sex-related monocyte activation differences between males and females with PD in bulk RNA-sequencing data of negatively-selected monocytes^31^; PD females were enriched for genes correlating with pro-inflammatory interferon-gamma expression programs, while males had more heterogenous alterations. No scRNA-Seq studies have yet addressed sex as a biological variable in peripheral immune cells. These recently published studies support the ability of single-cell transcriptomics to identify changes in immune cells in PD cohorts, and importantly, additional high-resolution datasets will improve statistical power, and thereby the opportunity to identify and understand PD risk factors.

To expand the data available for analyzing immune cell transcriptomic changes in PD, we analyzed freshly obtained PBMCs from a cohort of 12 PD patients and 12 healthy controls, equally distributed by sex, using scRNA-Seq. We identified differentially expressed genes and alterations in immune cell lineages in PD compared with control samples. Analysis of the dataset parsed by sex identified aspects of the PD-related changes that are, and are not, associated with sex. Independently analyzed data from these subsets identified additional pathways and gene expression profiles associated with PD. Trajectory analysis of the myeloid and T cell datasets indicated significant changes in cell distribution across the trajectory in PD compared with cells isolated from healthy control subjects. This work provides new evidence of peripheral immune cell changes in PD using high-resolution transcriptomics in a cohort designed to analyze sex as a variable.

## Methods

### Study Population

This study was approved by the Institutional Review Board of the Dartmouth Hitchcock Medical Center (Clinical Manifestations of Inflammasome Activity in PD, CPHS# 30209). Each participant provided informed consent, and all sampling and preparations were performed in accordance with Institutional and NIH guidelines. PBMCs were collected by clinical phlebotomists at Dartmouth-Hitchcock Medical Center and processed within 3 hours of collection. Purified PBMCs were either subjected to library preparation on the day of collection or preserved immediately using the 10x Flex gene expression assay, which was specifically designed to fix samples and preserve high-quality cells for scRNA-Seq analysis. Participants were selected for sequencing using deidentified medical records and self-reported patient survey data to identify comparable patients free from confounding comorbidities, including but not limited to autoimmune conditions and cancers. Patient population characteristics and covariate analysis are provided in Tables 1 and 2.

**Table 1:** Participant characteristics of single cell RNA-sequencing cohort collected at Dartmouth-Hitchcock Medical Center, Lebanon, NH from 2019-2023.

|  |  | **Controls**<br>*n* = 12 (%) | **PD**<br>*n* = 12 (%) | **Univariate *p*-value \*** |
| --- | --- | --- | --- | --- |
| **Sex** | **F** | 6 (50) | 6 (50) | **1** |
|  | **M** | 6 (50) | 6 (50) |  |
| **Age** | **Mean ± SD** | 61.5 ± 16.50 | 66.75 ± 6.50 | **0.323** |
| **Age category** | **<65** | 6 (50) | 4 (33.3) | **0.223** |
|  | **65–75** | 3 (25) | 7 (58.3) |  |
|  | **75+** | 3 (25) | 1 (8.3) |  |
| **Family history of PD** | **No** | 9 (75) | 8 (66.7) | **1** |
|  | **Yes** | 3 (25) | 4 (33.3) |  |
| **Disease duration** | **Mean ± SD** |  | 4.91 ± 4.61 | **0.080** |
|  | **F** |  | 7.17 ± 4.79 |  |
|  | **M** |  | 2.20 ± 2.77 |  |
| **Smoking** | **Never** | 9 (75) | 7 (58.3) | **0.665** |
|  | **Ever** | 3 (25) | 5 (41.7) |  |

\*calculated with chi-square or t-test

**Table 2:** Categorical medication use of single cell RNA-sequencing cohort.

|  |  | **Medication Category**<br>**n (%)** |  |  |  |  |  |  |  |  |  |
| --- | --- | --- | --- | --- | --- | --- | --- | --- | --- | --- | --- |
|  |  | **Neuro-modulators** |  | **Dopamine modulators** |  | **Anti-inflammatory** |  | **NSAID** |  | **Levodopa** |  |
| **Male** | **HC** | 1 | 17% | 0 | 0% | 3 | 50% | 2 | 33% | 0 | 0% |
| **Male** | **PD** | 4 | 67% | 3 | 50% | 3 | 50% | 3 | 50% | 2 | 33% |
| **Female** | **HC** | 3 | 50% | 1 | 17% | 1 | 17% | 1 | 17% | 0 | 0% |
| **Female** | **PD** | 4 | 67% | 2 | 33% | 3 | 50% | 3 | 50% | 6 | 100% |
| **Total** |  | 12 | 50% | 6 | 25% | 10 | 42% | 9 | 38% | 8 | 33% |

### Single cell RNA-sequencing (scRNA-Seq)

Single-cell RNA sequencing was performed using two comparable 10x Genomics workflows. Single-cell gene expression libraries were prepared using the Chromium Single Cell 3’ Reagent Kit v2 (10x Genomics) according to the manufacturer’s instructions (CG000052). Cells were loaded onto the Chromium Controller using the Chromium Single Cell A Chip Kit (PN-120236), and GEM generation, barcoding, and library construction were performed as specified in the manufacturer’s protocol. For preserved cell preparations, samples were processed using the Chromium Fixed RNA Profiling (Flex) Reagent Kit according to the manufacturer’s instructions (CG000580), using the Chromium G Chip Kit (PN-1000286) on the Chromium Controller. Samples were multiplexed using the Chromium Fixed RNA Profiling Probe Kit prior to library construction. Sequencing modalities are referred to herein as 3’ GEX and FLEX, respectively. All libraries were assessed for quality by Agilent TapeStation prior to sequencing. Libraries from both workflows were identically sequenced on an Illumina NextSeq 2000 targeting a minimum of 30,000 reads per cell using manufacturer-recommended read configurations (28 bp Read 1, 10 bp i7 index, 10 bp i5 index, 90 bp Read 2 for 3’ v2). Raw sequencing data were processed using Cell Ranger v8.0, each aligned to the mm10 reference genome using default parameters.

### Data Processing

The filtered barcode matrix outputs from CellRanger v8 were loaded into the R statistical programming environment (v 4.4.1) using the Seurat v5 R package and subjected to additional quality control filtering. Prior to filtering, there were nearly 220,000 cells. Cells exhibiting high proportions of reads mapping to the mitochondrial genome or low expression complexity were removed from downstream analysis (cells with a number of reads (nCount) <600, number of genes expressed (nFeature) <1000 and >5000, log10 genes per UMI <0.8, and mito-ratio of >20% were filtered out). Over 180,000 cells remained following the additional post-CellRanger quality control filtering. The mean number of reads per cell was 4,834 and the mean log10 genes per UMI was 0.93.

Expression of key cell-cycle regulators was identified using the cc.genes object available in Seurat, which contains cell cycle markers used to score cells for cell cycle stages. First, data were log-normalized using NormalizeData (Seurat). The most variable features were identified and scaled before statistically regressing out potential variation across the dataset attributable to cell-cycle phase (default nFeature of 2,000). A reciprocal PCA (RPCA)-based anchor method was used to integrate the dataset and perform batch correction. We applied the first 40 principal components to construct a shared-nearest neighbor graph and identify Louvain clusters and ultimately selected a resolution of 0.15 for initial analysis. We computed a uniform manifold approximation projection (UMAP) for visualization. Reference-based cell-type annotation was performed using the Azimuth PBMC atlas^32^. The atlas was loaded into R and identities were set to a resolution of “celltype.l1” to include monocytes, CD4+ T cells, CD8+ T cells, Natural Killer cells (NK), B cells, and dendritic cells (DC). We identified transfer anchors by querying the integrated dataset against the reference atlas via FindTransferAnchors, using the first 30 PCA dimensions for dimensionality reduction. The cell type identities were transferred to the dataset using TransferData (Azimuth), maintaining all annotations, and the associated prediction scores and metadata were added via AddMetaData. Cells with low-quality predictions scores (<0.5) were removed.

### Differential Gene Expression and Ontology Analysis

To identify changes in gene expression, data layers (Seurat v5) were joined, and the FindMarkers function was used to identify differentially expressed genes (DEGs) between PD and control per cell type identity (logfc.threshold=0.1, test=MAST). We performed differential gene expression analysis using the Model-based Analysis of Single-cell Transcriptomics (MAST). MAST explicitly models zero-inflation and cell-to-cell variability in scRNA-Seq to improve sensitivity and power in detecting more subtle transcriptional changes in heterogeneous cell populations. To determine cell-type-specific and shared DEGs, we used UpSetR for visualization of these gene sets^33^. MA plots were created by calculating log2-transformed M-values (mean percent expression) and the log2 fold change. For p-values associated with DEGs exceeding the lower limit of extremely small p-values (R limit), the adjusted p-values at “0” were temporarily assigned values below the limit (1e-320) to allow for visualization in certain plots (e.g., MA). To investigate if mitochondrial genes that appeared in differential gene expression analyses between PD males and PD females, we reperformed the MAST framework with mitochondrial read fraction and library size included as covariates to control for potential mitochondrial content-driven artifacts. Inclusion of these covariates did not account for the observed mitochondrial gene dysregulation, indicating these effects were not attributable to mitochondrial content-related technical artifact.

We performed gene ontology (GO) enrichment analysis using enrichGO (clusterProfiler)^34^. DEGs were stratified by cell type and direction of regulation (upregulated or downregulated in PD), with Bonferroni adjusted p-values <0.05 and log2 fold changes>0.1 and <-0.1, respectively. Enrichment was conducted for the Biological Process (BP) ontology using the org.Hs.eg.db database for gene annotation. Dysregulated gene pathways were identified using all statistically significant DEGs, regardless of fold-change directionality. Resulting matrices were visualized using ComplexHeatmap^35^. Sex-specific analyses were performed as described above for the full dataset. Parkinson’s disease-associated genes discussed in the Results section were identified using the Protein ANalysis THrough Evolutionary Relationships (PANTHER) Parkinson’s disease pathway group^36^.

### Cell subtype analyses

To perform a deeper characterization of myeloid cells and T cells, the raw data were processed again after subsetting the dataset to include only those cells assigned to either of these cell types in the initial analyses. These data were subject to the same quality control preprocessing, but the reference-based cell type identification was performed prior to integration at a higher resolution (“celltype.l2”), allowing us to assign the following lineage subtype labels: myeloid (CD16+ monocytes, CD14+ monocytes, cDC2, pDC, and ASDC) and T cells (CD4+ Naïve, CD4+ Tcm, CD4+ Tem, CD4+ CTL, CD4+ Proliferating, CD8+ Naïve, CD8+ Tcm, CD8+ Tem, CD8+ Proliferating, Treg, MAIT, dnT, and gdT). These datasets were normalized via SCTransform (Seurat)^37^, then integrated using the same anchor-based RPCA approach described above. Differential gene expression and gene ontology analyses were performed independently on these datasets as described above.

### Trajectory Analysis

To investigate transcriptional dynamics within the myeloid cell and T cell subsets described above, we performed trajectory analysis using Monocle3^38^. Count matrices from each sample layer were extracted and merged. These data were assembled into a CellDataSet (CDS) (SeuratWrappers), and the original subset’s UMAP coordinates were transferred to the CDS. Dimensionality reduction and clustering were performed using preprocess_cds and cluster_cells (Monocle3). The trajectory graph was made using the learn_graph function. We assigned root clusters to each object based on gene expression. For T cells, we selected root cells from the cluster with the highest expression of naïve-like T cell development markers (*CCR7*, *SELL/CD62L*, *TCF1/TCF7*, and *LEF1*)^39,40^, and no activation or exhaustion markers (*IFNG*, *GZMB, PD*-*1*, *CTLA4*)^41–43^. For myeloid cells, we selected root cells with the highest expression of genes associated with emergence from the bone marrow in CD14+ monocytes (*CCR2, CXCL4*)^44–46^. We ordered the cells in pseudotime from this assigned root cluster. Pseudotime trajectory values were assigned to the original Seurat object for visualization. Comparisons between trajectory relationships between PD and control were performed using Wilcoxon rank-sum tests.

## Results

### Patient Population

Single-cell RNA sequencing was used to profile PBMCs isolated from blood samples obtained from a cohort of 12 PD patients and 12 healthy control subjects, evenly distributed in both groups by sex (Fig. 1a). Table 1 summarizes the demographic characteristics of the selected study population. There were no significant differences in mean age or age distribution across sampled ages between controls and PD, combined or by sex. There were also no significant differences in self-reported family history of PD or smoking behavior identified, as assessed by chi-square analysis or t-test. The average disease duration was 4.91±4.61 years for all PD participants, 2.20±2.77 years for males and 7.17±4.79 years for females and was not significantly different between groups. Table 2 quantifies various groups of medications used by participants as recorded in their deidentified medical records. Medications are an important covariate to consider, as certain medications such as non-steroidal anti-inflammatory drugs (NSAID) and dopamine-replacement therapies interact with the immune compartment.

**Figure 1:**
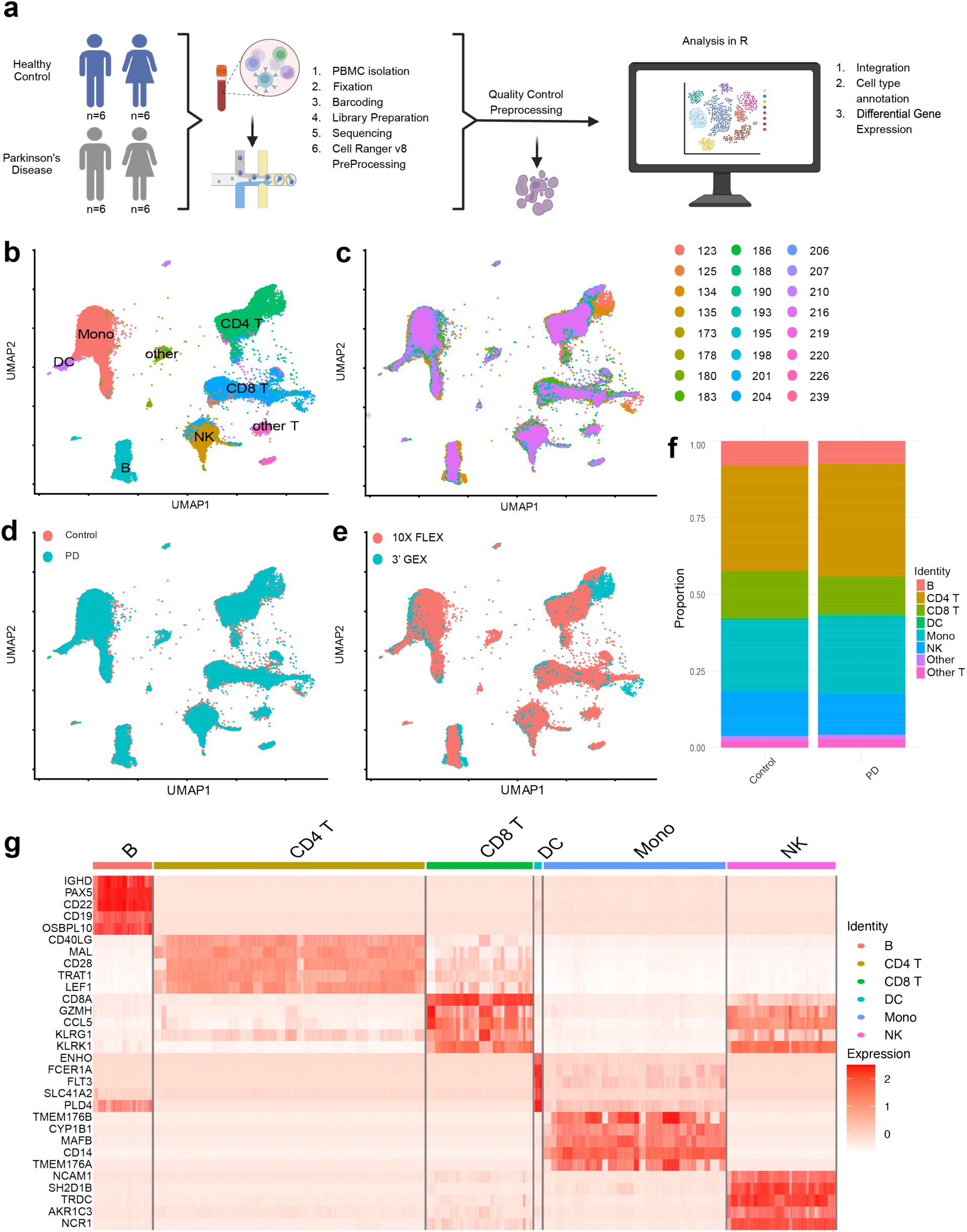
Single cell transcriptomic analysis of PBMCs identify major immune cell types in PD and controls. **a)** Schematic of experimental overview. **b to e)** UMAP of >100,000 PBMCs from participant cohort colored by cell types: monocytes (Mono), dendritic cells (DC), CD4+ T cells (CD4 T), CD8+ T cells (CD8 T), natural killer cells (NK), B cells, other T cells subsets (other T), and other cells (**b**); colored by participant deidentified study ID (**c**); colored by disease status (HC or PD) (**d**); and colored by 10x Genomics sequencing modality (3’ GEX or FLEX) (**e**). **f**) PBMC cell type proportions by disease status (HC or PD). **g)** Heatmap of top marker genes per cell type. n=24 participants.

### Parkinson’s disease-specific transcriptomic signatures in PBMCs

Unsupervised clustering identified distinct clusters that were assigned to specific cell types through reference-based annotation (Fig. 1b). Louvain algorithm clustering at resolution of 0.15 resulted in 22 clusters (Supplementary Fig. 1, Supplementary Table 1). We verified that these cluster identities included myeloid cells (clusters 1, 6, 7, 11, 20) with canonical markers *CSF1R, LST1,* and *CD163*; CD4+ T cells (clusters 0, 5, 9, 17) with *CD40LG, FOXP3,* and *MAL*; CD8+ T cells (clusters 3, 8, 13) with *CD8A* and *EOMES*; natural killer (NK) cells (clusters 2, 12, 14) with *NCR1, NCAM/CD56,* and *GZMB*; and B cells (clusters 4, 15, 18, 19) with *JCHAIN, IGHD/IGHA1*, and *CD22*. The remaining cluster 10 was characterized as “other T cells,” and clusters 16 and 21 were characterized as “unknown” and not included in subsequent analysis. Immune cell populations were comparable across PD and control patients (Fig. 1b-e). Quantitative analysis of cell type proportions suggests subtle shifts in myeloid and T cell type proportions in PD as previously reported^20,47,48^, but these shifts, when analyzed as percentage of total cells adjusted per participant, do not reach statistical significance in our dataset (Fig. 1f, Supplemental Table 2). Unsupervised analysis identified canonical markers of key lineages, such as CD14 for monocytes, CD8a for CD8+ T cells, and CD19 in B cells (Fig. 1g).

Differential gene expression analysis across cell populations revealed both cell-type-specific and shared transcriptional alterations associated with PD (Fig. 2a-b, Supplemental Tables 3-8). The UpSet plot reveals that CD8+ T cells harbor the most differentially expressed genes in our PD cohort, followed by monocytes and B cells. DEGs shared by CD8+ T cells and monocytes was the largest combined gene set. Gene ontology analysis of all dysregulated genes, upregulated genes, and downregulated genes reveals widespread reprogramming unique to each cell type (Fig 2c-e). The data include multiple pathways related to respiration and metabolic regulations consistent with prior reports indicating changes in immune cell activation state in PD^49–52^.

**Figure 2:**
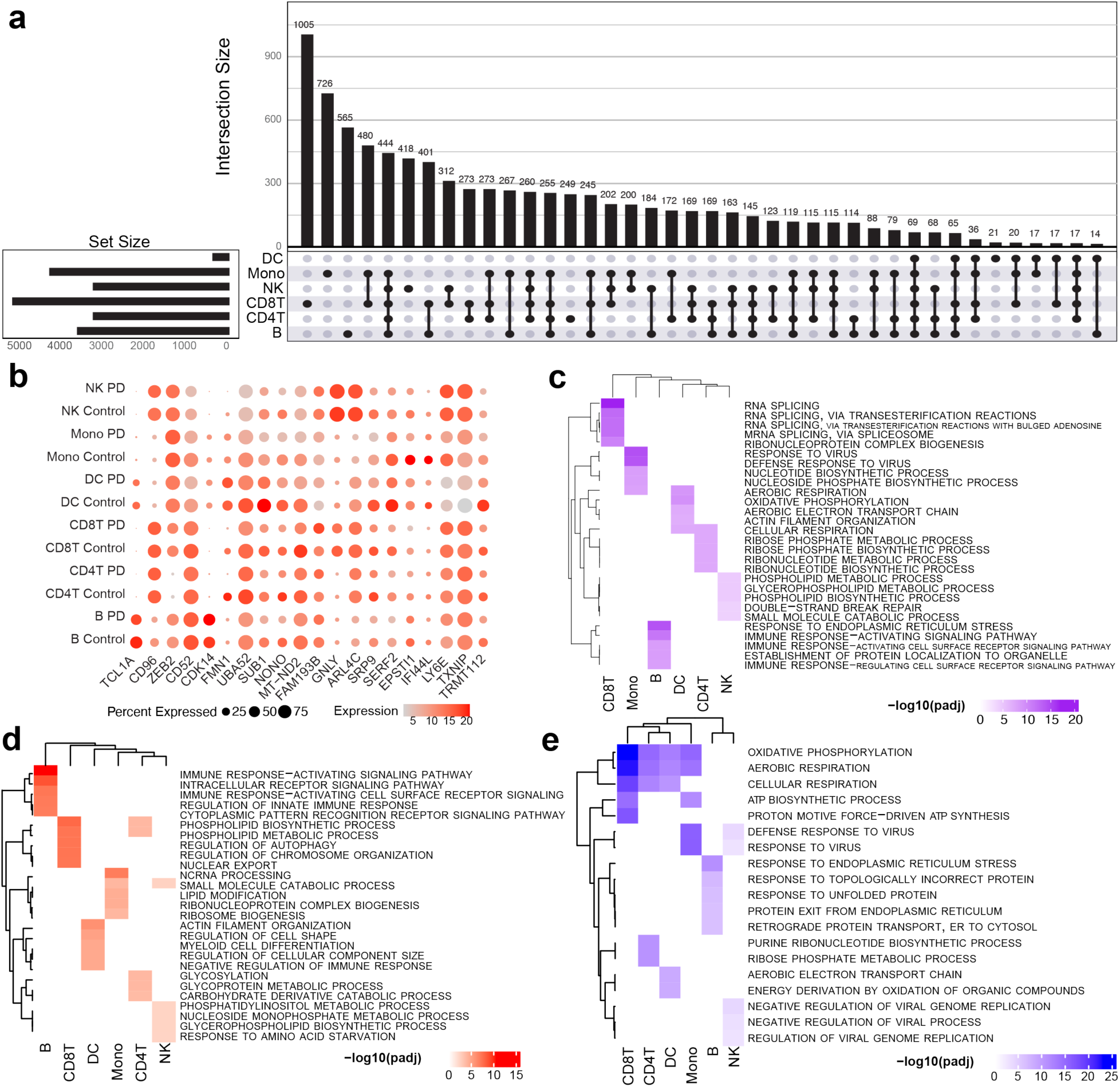
Differential gene expression analysis reveals PD disease-state transcriptional programming alterations across cell types. **a)** UpSet plot of differentially expressed gene (DEG) sets in PD that are unique in 6 cell types and sets in common have two or more cell types. **b)** Dot plot of top DEGs per cell type. Shared genes are represented once. Dot size represents percent of cells the gene is expressed in, and color represents scaled and normalized average expression. **c-e)** Heatmaps show top 5 gene ontology pathways of all significant DEGs per cell type (**c**, purple), only enriched genes (**d**, red), and only depleted genes (**e**, blue). For all panels, DEGs are included with an Bonferroni-adjusted p-value <0.05.

### Parkinson’s disease-specific transcriptomic signatures in males and females

Differential gene expression analysis identified sex-specific transcriptional alterations in the major immune cell types in PD (Fig. 3). Cell type proportions split by disease-state indicates minimal proportional changes in cell types between males and females in PD and control groups that do not meet statistical significance when accounting for sex (Fig. 3a-b, Supplemental Table 9). Dot plots of DEGs per cell type in PD females compared to PD males show significant changes in several mitochondrial genes such as *MT-ND4, MT-ND4L, MT-ND6,* which may be reflective of differential bioenergetic programming between males and females in PD^53^ (Fig 3c). *IFITM2, ZFP36L2,* and *JUNB* are DEGs that further demonstrate fine-tuning of the immune functions of these cells as they are involved in inflammatory signaling and responses^54^. We conducted ontological analysis of the top five differentially expressed pathways per cell type in both positive and negative directions between male PD and female PD (Fig. 3d), only enriched genes (Fig. 3e), and only depleted genes (Fig. 3f). Overall, the DEG analysis identified sex-specific transcriptional alterations in all the major immune cell types, suggestive of sex-related immune traits associated with PD.

**Figure 3:**
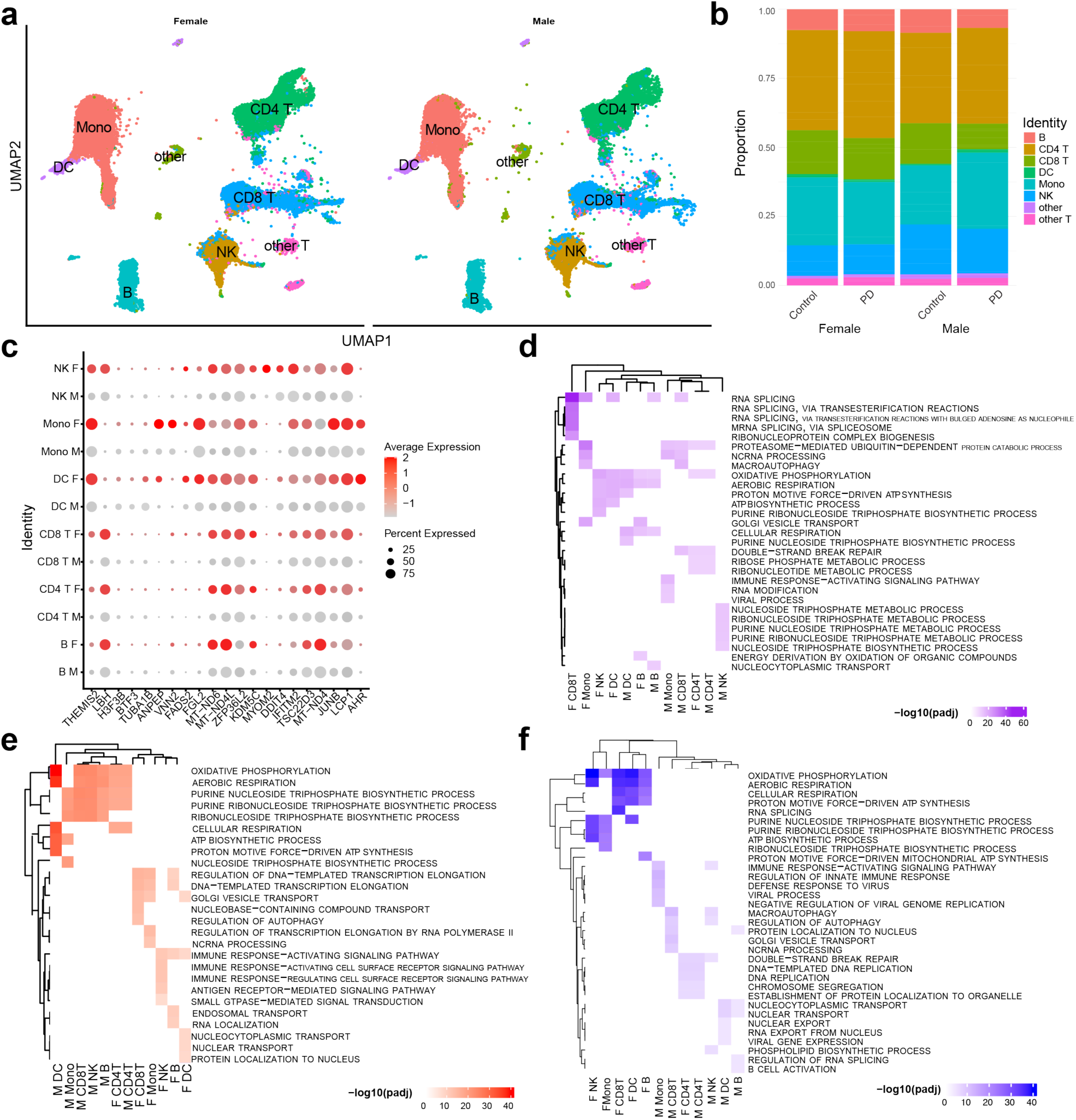
Differential gene expression analysis reveals interaction of sex in PD disease-state transcriptional programming alterations across cell types. **a)** UMAP of all subjects split by sex. **b)** Stacked bar plot of cell type proportions split by disease-state and sex. **c)** Dot plot of top DEGs in PD females compared to males. Shared genes are represented once. Dot size represents percent of cells gene is expressed in, and color represents scaled and normalized average expression. **d-f)** Heatmaps show top 5 gene ontology pathways of all significant DEGs per cell type between male PD and female PD (**d**, purple), only enriched genes (**e**, red), and only depleted genes (**f**, blue). Pathways in male cell types are denoted with M and females with F. For all panels, DEGs are included with an Bonferroni-adjusted p-value <0.05.

### Sub-clustering reveals additional changes in the myeloid lineage

Re-analysis of parsed mononuclear myeloid cells (see *Methods*) identified five cell types: two corresponding to monocytes and three corresponding to dendritic cells. Azimuth cell type identification confirmed that these populations represented CD14+ monocytes (*classical monocytes*), CD16+ monocytes (*non-classical and intermediate monocytes*), conventional dendritic cells (cDC2), plasmacytoid dendritic cells (pDC), and AXL+SIGLEC6+ dendritic cells (ASDC) (Fig. 4a). CD14+ monocytes were consistently the most abundant, followed by CD16+ monocytes (Fig. 4b). We again observed no statistically significant differences in overall proportions of immune lineages between PD and healthy controls (see also Fig. 1f), possibly due to variability between participants (Fig. 4b, Supplemental Table 10).

**Figure 4:**
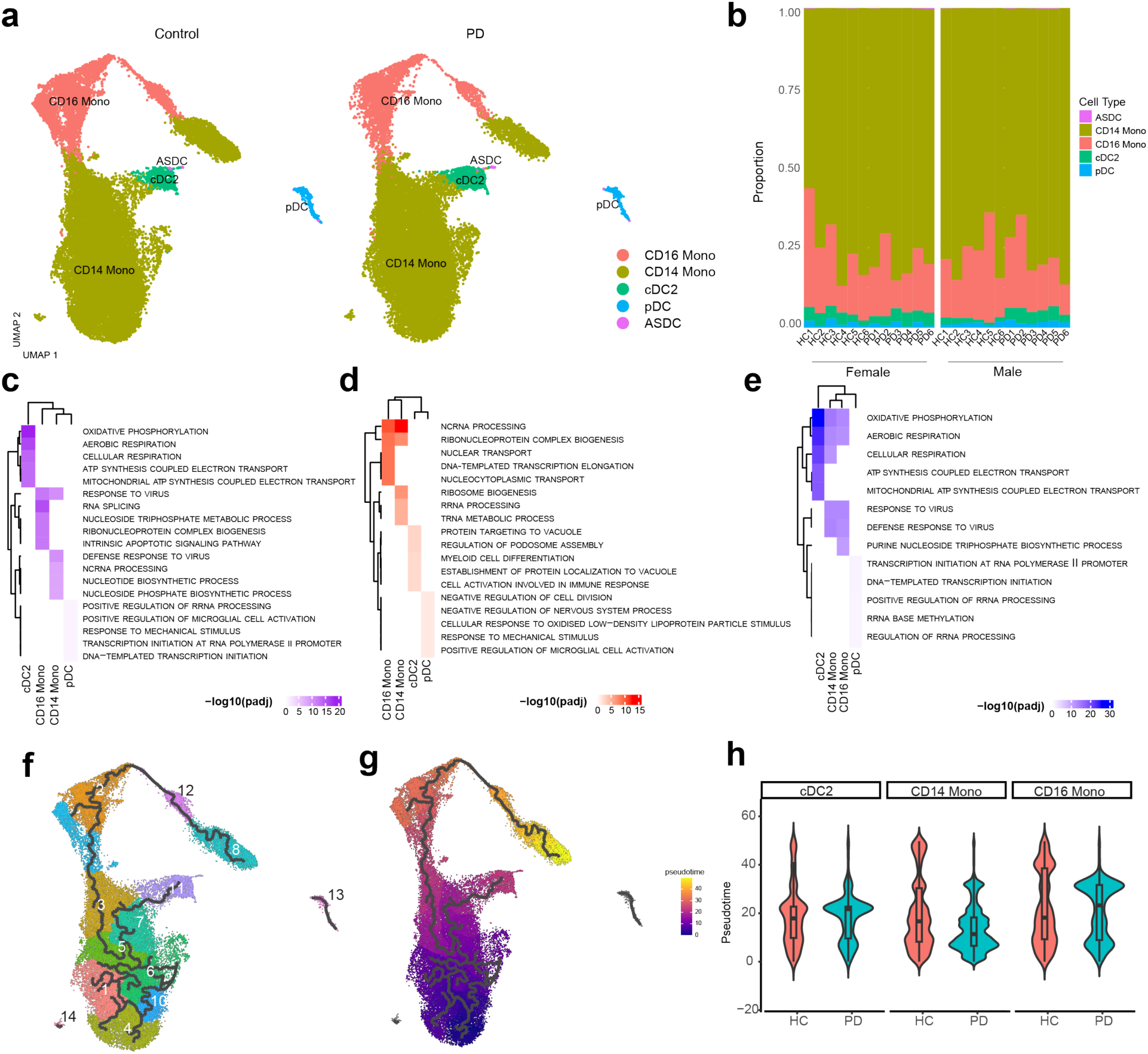
Deeper characterization of myeloid cell subtypes. **a)** UMAP of all subjects split by disease state and clustered by cell-type identity. **b)** Stacked bar plot of cell type proportions of myeloid cell subsets per subject. **c-e)** Heatmaps show top 5 gene ontology pathways of all significant DEGs per cell type (**c**, purple), only enriched genes in PD (**d**, red), and only depleted genes in PD (**e**, blue). For all panels, DEGs are included with an FDR-adjusted p-value <0.05**. f-g**) Trajectory analysis of myeloid cells clustered by pseudo-temporal changes (**f**) and colored by pseudotime (**g**). Black lines denote trajectories beginning at the assigned node (cluster 4) (pseudotime=0). **h)** Violin plots of cell densities across the trajectory (pseudotime, y-axis) per cell type between disease-states. Distributions are significantly different between groups in cDC2 between pseudotime score 5-10, 30-35, and 35-40; in CD14 monocytes between 5-15, 20-40; and in CD16 monocytes between 5-10, 20-25, and 30-40.

Differential gene expression and ontological analysis on each of the five myeloid subsets identified a significant downregulation of oxidative phosphorylation-related genes in cDC2, CD14+ monocytes, and CD16+ monocytes (Fig. 4c-e). In both CD14+ and CD16+ monocytes, genes involved in transcription and protein biosynthesis-related pathways are upregulated, such as transcription elongation, ribosome biogenesis, and nucleocytoplasmic transport (Fig. 4d). cDC2 cells show significant upregulation of genes associated with myeloid cell differentiation and cell activation in immune response (Fig. 4d). Pathways including aerobic respiration and cellular respiration are also significantly downregulated across these cell types (Fig. 4e). In cDC2, metabolism-related pathways, ATP synthesis coupled electron transport, and mitochondrial ATP synthesis coupled electron transport are also downregulated, demonstrating a major shift in cDC2 metabolic programming (Fig. 4e). Genes related to defense against virus and response to virus that are typically associated with type I interferon signaling are downregulated in monocytes (Fig. 4e).

To compare the distributions of myeloid phenotypic states between PD and control subjects, we performed trajectory analysis on sub-clustered myeloid cells. We overlaid the pseudotemporal trajectory onto the original dimensionality coordinates and “colored” the cells by pseudotime (Fig. 4f-g). We defined a node representing the gene expression profile of monocytes immediately following emergence from bone marrow (see *Methods*) (Fig. 4g). This node was assigned to these cells to begin the trajectory and is observable as cluster 4 (Fig. 4f), comprised of CD14+ cells that express high levels of *CCR2, CD14*, and monocyte-stemness markers like *CD36* and *CD64.* The distributions of these cells along the trajectory are represented by violin plots, and these plots demonstrate differences in cell numbers along the trajectory continuum (Fig. 4h). The distributions are significantly different between groups in cDC2 between pseudotime score 5-10, 30-35, and 35-40; in CD14 monocytes between 5-15, 20-40; and in CD16 monocytes between 5-10, 20-25, and 30-40. Subset characterization of myeloid cells and trajectory analyses reveal nuanced changes in transcriptional signatures and differences in the densities of cell sub-populations in PD.

To explore the potential of sex-specific differences in PD-related myeloid gene expression profiles, we performed additional analysis on the two most abundant myeloid subsets, CD14+ and CD16+ monocytes. DEG analysis revealed the vast majority of DEGs in CD14+ and CD16+ monocytes are shared between males and females (Fig. 5a, Fig 8). Male CD14+ monocytes show the largest gene set of unique DEGs, followed by male CD16+ monocytes (Fig. 5a). When compared to controls, male CD14+ monocytes upregulate genes associated with metabolic activation (*MT-ND1/2/3/4, NDUFS5, UQCR11, COX5B, ATP5MPL*), protein homeostasis, and cellular machinery (*PRKN, BTF3, LSM3, TMA7, MYL12B, ARPC3, H3F3B, SPARC*) (Fig. 5b).

**Figure 5:**
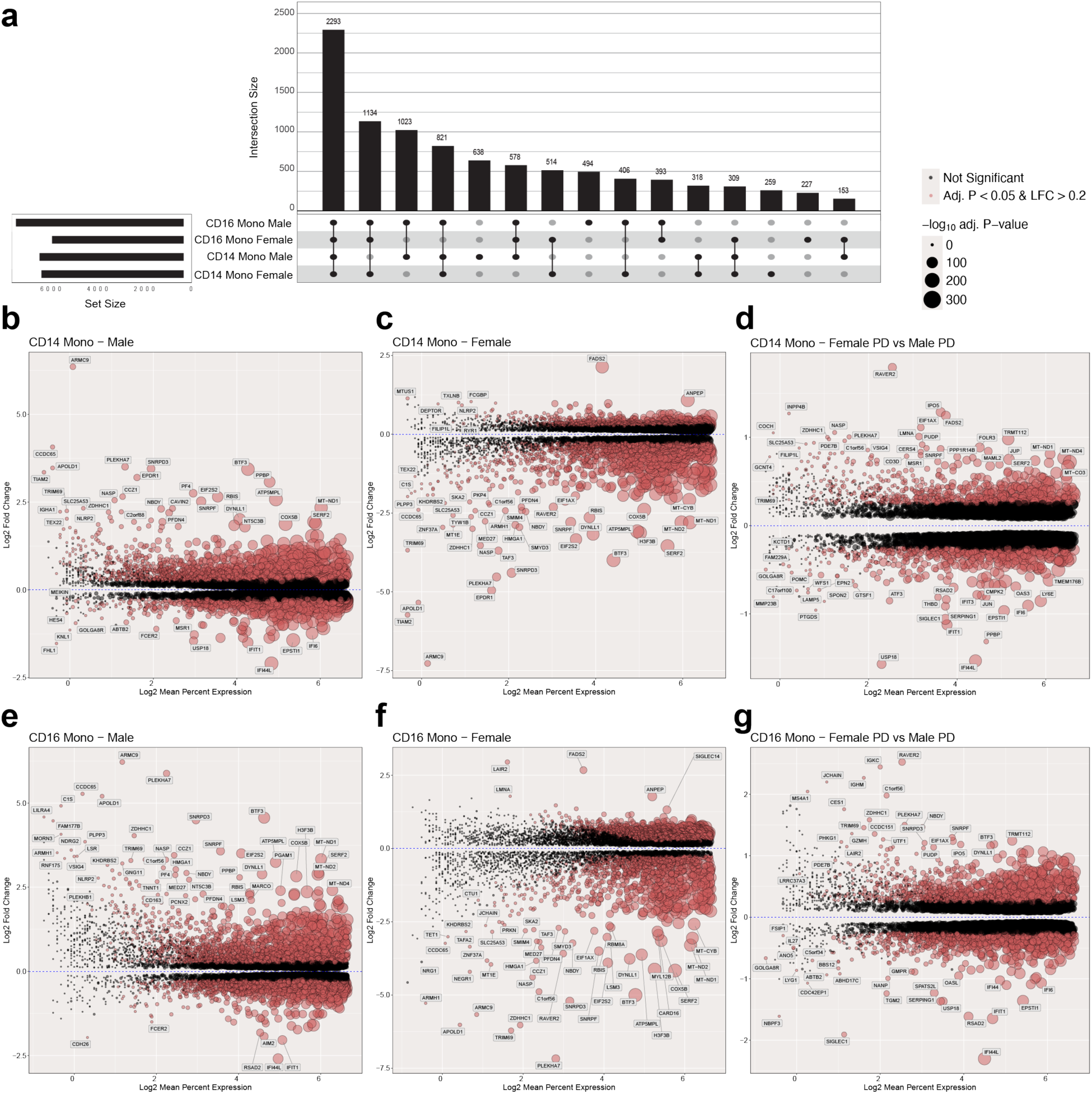
Transcriptomic comparison of differentially expressed genes (DEGs) across sex and disease status in CD14+ and CD16+ monocytes. **a**) UpSet plot of DEG sets in PD that are unique in 2 cell types in males and females and sets in common have two or more cell sets. **b-d**) MA plots show DEGs in CD14+ monocytes in **b**) males **c**) females and **d**) PD females vs. PD males. Genes above the x-axis are enriched in PD males and PD females compared to control males and control females, respectively (**b-c**). Positive Log2 Fold Change indicates genes enriched in PD females and negative Log2 Fold Change indicates genes enriched in male PD. (**d**). **e-g**) MA plots show DEGs in CD16+ monocytes in **e**) males **f**) females and **g**) PD females vs. PD males. Positive Log2 Fold Change indicates genes enriched in PD females and negative Log2 Fold Change indicates genes enriched in male PD. Statistically significant genes are colored red, and the size of the circle corresponds to -log10(Bonferroni-adjusted p-value).

Male PD CD14+ monocytes downregulate genes associated with type I interferon response and immune regulation (*IFIT1/2/3/5, IFI44/44L, MX1/2, OAS1/2/3, RSAD2, IRF7, BCL3, SP140L, HSH2D, VSTM1, CD52, LGALS3BP*) (Fig. 5b). PD female CD14+ monocytes upregulate genes associated with activation, cytokine signaling, and inflammation response (*ANPEP*, *CLEC10A* (*CD301*), *IRF4, LYZ, MRC1* (*CD206*), *TLR2/4, NLRP2, FADS2*); oxidative stress (*HSPA1B*, *NFE2L3, AHR, NQO2, DUSP6/7/22*); and cell adhesion and migration (*ARPIN, GSN, BAIAP2, CCR2, CCR5, SELL* (*CD62L*), *ICAM1, ITGAL, ITGB2*) (Fig. 5c). Female CD14+ monocytes downregulate transcription and translation associated genes (*BTF3, ZBTB20, MBD5*) (Fig. 5c). When we compare PD female CD14+ monocytes to PD male CD14+ monocytes, female cells are enriched for immune activation genes (*GZMH, VSIG4, FCGBP, FOLR3*), while male cells are enriched for inflammation response (*CCL3, RGS1, SIGLEC1, EPSTI1, SERPING1, PTGDS, IFI44/IFI44L, IFIT1/3, RSAD2, OAS3, LY6E*) and stress response/apoptosis related genes (*ATF3/6B/7, JUN, JDP2, TUBA1B*) (Fig. 5d). When compared to controls, male CD16+ monocytes similarly upregulate tumor necrosis factor (TNF) family and innate immune activation related genes (*CD14, TYROBP, CD36, FCGR1A, C1S*); vascular interaction/cell migration related genes (*PLEKHA7, CCR2, CD47, AGER, JAM3*), and oxidative stress genes (*PPIA, PSMA7, PSMB7, FOS, PRDX2*); they downregulate type I and type II interferon associated genes (*IFI44L, IFI6, IFIT1/2/3, IFITM1, OASL, EPSTI1, AIM2, MEFV, IRF2/7, STAT1, CXCL16*) (Fig. 5e). PD female CD16+ monocytes upregulate genes associated with immune regulation and inflammation (*TNF, VNN1, NLRP2, LAIR2, SIGLEC14, PELI3*) and lipid metabolism (*FADS2, CYP4F22, ANPEP*). CD16+ female monocytes genes associated with protein aggregation and stress response (*SERF2, EIF2S2, SNRPF, SNRPD3*); nuclear and chromatin organization (*H3F3B, HMGA1, NASP, ZNF37A*); and Parkinson’s disease-related genes (*YWHAB, FGR, PSMA7, PSMB3, MAPK1, HSPA6, GRK2*) (see Methods) (Fig. 5f). When we compare PD female CD16+ monocytes to PD male CD16+ monocytes, female cells are enriched for RNA modification-associated genes (*SNRPD2, SNRPF, TRMT112*), while male cells are enriched for genes associated with innate defense response (*OAS3, TRIM22, CLEC12A, EIF2AK2*) (Fig. 5g).

### Sub-clustering reveals additional changes in the lymphoid lineages

The strategy described above to analyze myeloid lineages was applied to T cells, identifying thirteen T-cell subtypes: CD4+ Naïve, CD4+ CTL (Cytotoxic), CD4+ Tcm (central memory), CD4+ Tem (effector memory), CD4+ Proliferating, CD8+ Naïve, CD8+ Tcm, CD8+ Tem, CD8+ Proliferating, Treg (regulatory T cell), gdT (γδ T cell), dnT (double negative T cell), and MAIT (Mucosal-associated invariant T cell) (Fig. 6a). There were no statistically significant changes in T cell subtype proportions between PD and controls, and there is noticeable heterogeneity in the T cell compartment fractions (Fig. 6b, Supplemental Table 10). CD4+ Tcm and CD8+ Tem were among the most abundant and therefore subject to further analysis. DEG analysis was performed on the CD4+ Tcm and CD8+ Tem subsets, comparing PD to healthy subjects. PD-related changes in CD4+ Tcm included those categorized as cellular respiration and related biological processes, as well as regulation of autophagy and RNA splicing (Fig. 6c). CD4+ Tcm upregulated pathways included aerobic and cellular respiration and oxidative phosphorylation (Fig. 6d). Downregulated pathways in CD4+ Tcm included lipid metabolism and biosynthesis, transcription, and TORC1 signaling (Fig. 6e). Ontologic analysis of PD CD8+ Tem cells identified alterations in RNA splicing, macroautophagy, and proteasome activity related pathways compared to cells from healthy individuals (Fig. 6c). These cells upregulate aerobic respiration, oxidative phosphorylation (OXPHOS), and cellular respiration, while pathways for regulation of autophagy and transcription are downregulated (Fig. 6d-e). These findings indicate changes in abundant T cell populations associated with PD broadly indicative of enhanced metabolic activity.

**Figure 6:**
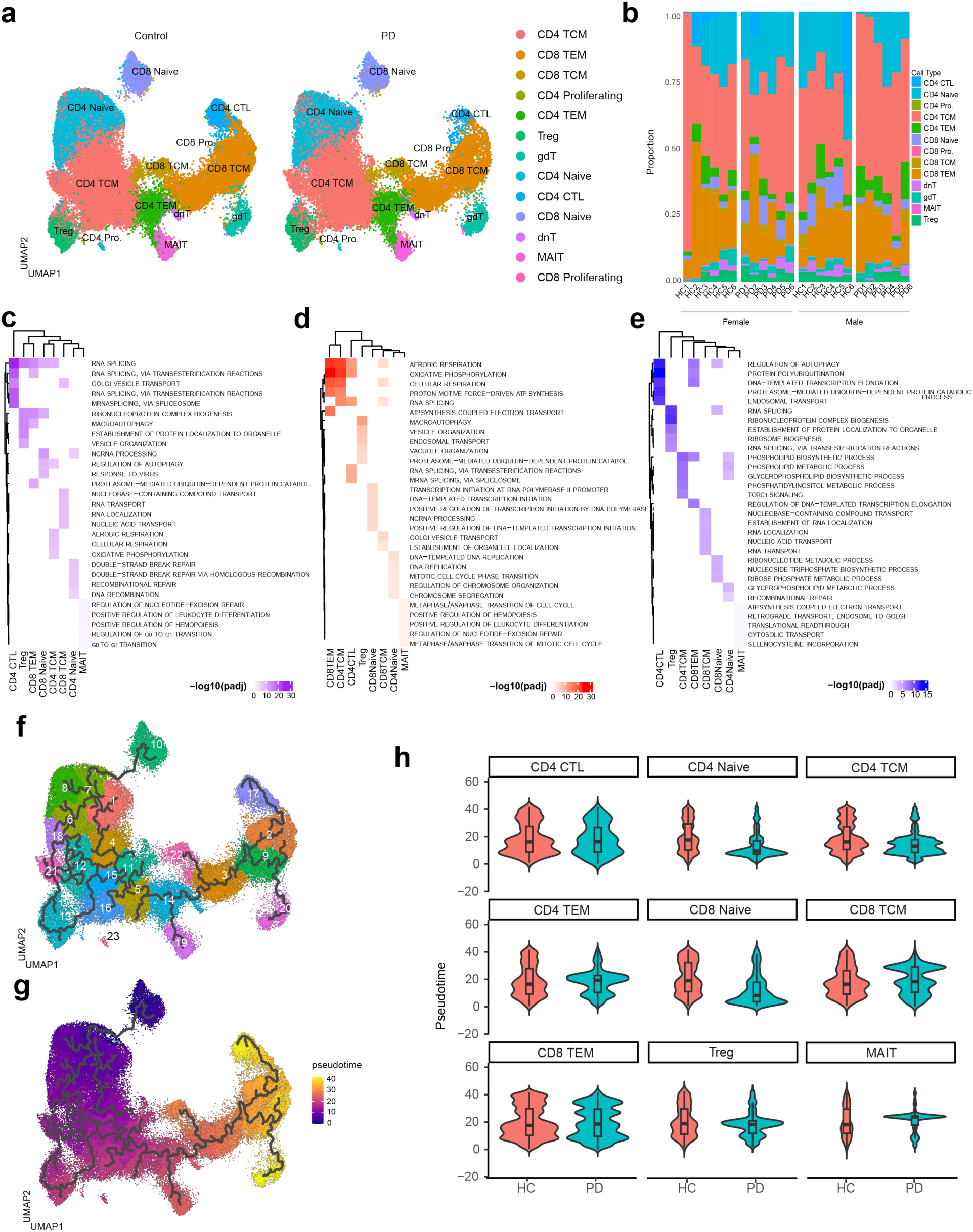
Deeper characterization of T cell subtypes. **a**) UMAP of all subjects split by disease state and clustered by cell-type identity. **b**) Stacked bar plot of cell type proportions of T cell subsets per subject. **c-e**) Heatmaps showing gene ontology pathways of all significant DEGs per cell type (**c**, purple), only enriched genes in PD (**d**, blue), and only depleted genes in PD (**e**, red).

We performed trajectory analysis as described above to investigate the dynamics of these T cell states between PD and control samples (Fig. 6f-h). In this case, we identified cluster 10 as the root node based on the expression of genes associated with naïve (versus exhausted) cells (see *Methods*). We adopted this strategy because unlike circulating myeloid cells, which share the CD14+ monocyte progenitor, the common progenitors of CD4+ and CD8+ T cells are not circulating in the blood. In this regard, naïve versus exhausted trajectories better reflect differences in transcriptional programming between PD and healthy controls as opposed to T cell development from progenitors. The distributions of these T cells along these trajectories are represented by violin plots, and these plots demonstrate differences in cell densities along the trajectory continuum (Fig. 6h). The distributions are significantly different between groups in CD4+ CTL at pseudotime score 30-40; in CD4+ naïve between 0-40; in CD4+ Tcm between 0-30; in CD4+ Tem between 20-30; in CD8+ naïve between 0-10, 20-30; in CD8+ Tcm between 20-30; in CD8+ Tem between 20-40; in Treg between 10-30; in MAIT between 20-30. These differences point to distributional changes of transcriptional programming between healthy controls and PD, and they provide an opportunity to understand minute changes in the transcriptome and abundance of cells in similar transcriptomic states.

For all panels, DEGs are included with a Bonferroni-adjusted p-value <0.05. **f-g**) Trajectory analysis of T cells clustered by trajectory changes (**f**) and colored by pseudotime (**g**). Black lines denote trajectories beginning at the node (cluster 10) (pseudotime=0). **h**) Violin plots of cell number densities across pseudotime (y-axis) per cell type between disease-states. Distributions are significantly different between groups in CD4+ CTL at pseudotime score 30-40; in CD4+ naïve between 0-40; in CD4+ Tcm between 0-30; in CD4+ Tem between 20-30; in CD8+ naïve between 0-10, 20-30; in CD8+ Tcm between 20-30; in CD8+ Tem between 20-40; in Treg between 10-30; in MAIT between 20-30.

We analyzed sex differences for the two most abundant lymphoid subsets, CD8+ Tem and CD4+ Tcm. The vast majority of DEGs in CD8+ Tem and CD4+ Tcm are shared between males and females (Fig. 7a, Fig. 8). Male CD8+ Tem show the largest gene set of unique DEGs, followed by male CD8+ Tem (Fig. 7a). When compared to controls, male CD8+ Tem upregulate genes associated with metabolism (*ATP5MPL, COX5B, PGAM1, FTH1*), cellular stress, and immune regulation (*SOD1, PPIA, PRDX1, CCRL2, CXCR5, SERF2*) (Fig. 7b). These male cells downregulate genes related to cytotoxic capacity, cytokine signaling, and activation (*FASLG, CD226, PRSS23, THEMIS2, CX3CR1, SOCS7, CXCR1/2, GRN, LAMP1, CX3CR1, ADAM10*), as well as those related to adhesion and motility (*ITGAM/CD11b, ITGB1/CD29, ADGRG1, S1PR5, ITGAL, CX3CR1, PLEKHG3*) (Fig. 7b). PD female CD8+ Tem upregulate lipid metabolism and stress related genes (*FADS2, ABCG1, AMPD3, FKBP5*), as well as inhibitory signaling genes (*LAIR1, TNFRSF25, THEMIS2, UBASH3B*) (Fig. 7c). Female PD CD8+ T cells downregulate genes associated with activation and cytotoxicity compared with healthy control females (*CCL3, TYROBP, DYNLL1*) (Fig. 7c). When we compare PD female CD8+ Tem to PD male CD8+ Tem, female cells are enriched for genes associated with effector activation compared to PD males (*IFNG, CCL3, DAPK2, BCL7A, TYROBP*), while male PD CD8+ Tem are enriched for genes that are responsive to type I and type II interferons (*IFI44L/IF44, IFI6, IFIT3, EPSTI1, CCR2, SLC7A5, TXK*) (Fig. 7d). PD male CD4+ Tcm upregulate oxidative phosphorylation and respiration associated genes (*ATP5PF, COX7C, COX5B, ATP5F1E, MT-ND1/2*) and Parkinson’s disease-related proteosome genes (*PSMB7, PSMB3, PSMA5*) relative to controls (Fig 7e). These cells downregulate genes associated with inflammatory cytokine response (*RGS1, ITGB1/CD29, NRAS, JUNB, BCL3, IL2RB*) (Fig. 7e). PD female CD4+ Tcm upregulate lipid metabolism genes (*FADS2, LPAR6, SLC30A1*) and activation regulatory/modulatory genes (*PLXNA4, ADAM23, RAB11FIP5, SNX25, THEMIS2, PILRB*), while they downregulate genes associated with oxidative phosphorylation/ATP synthesis (*MT-ND1, MT-ND2, COX5B, UQCRH, ATP5PF, ATP5MPL*) and RNA/protein processing (*PSMB3, PSME2, DAD1, EIF1AX, TMA7, SRSF10, LSM3, PPIA, SERF2*) (Fig 7f). PD female CD4+ Tcm are enriched for defense and stress response genes (*ALOX5, TMEM258, NCR3, PSMB3*) compared to PD males, while PD males are enriched for genes associated with inflammatory cytokine response (*CCR2, JUNB, BRAF, SMAD7*) (Fig. 7g).

**Figure 7:**
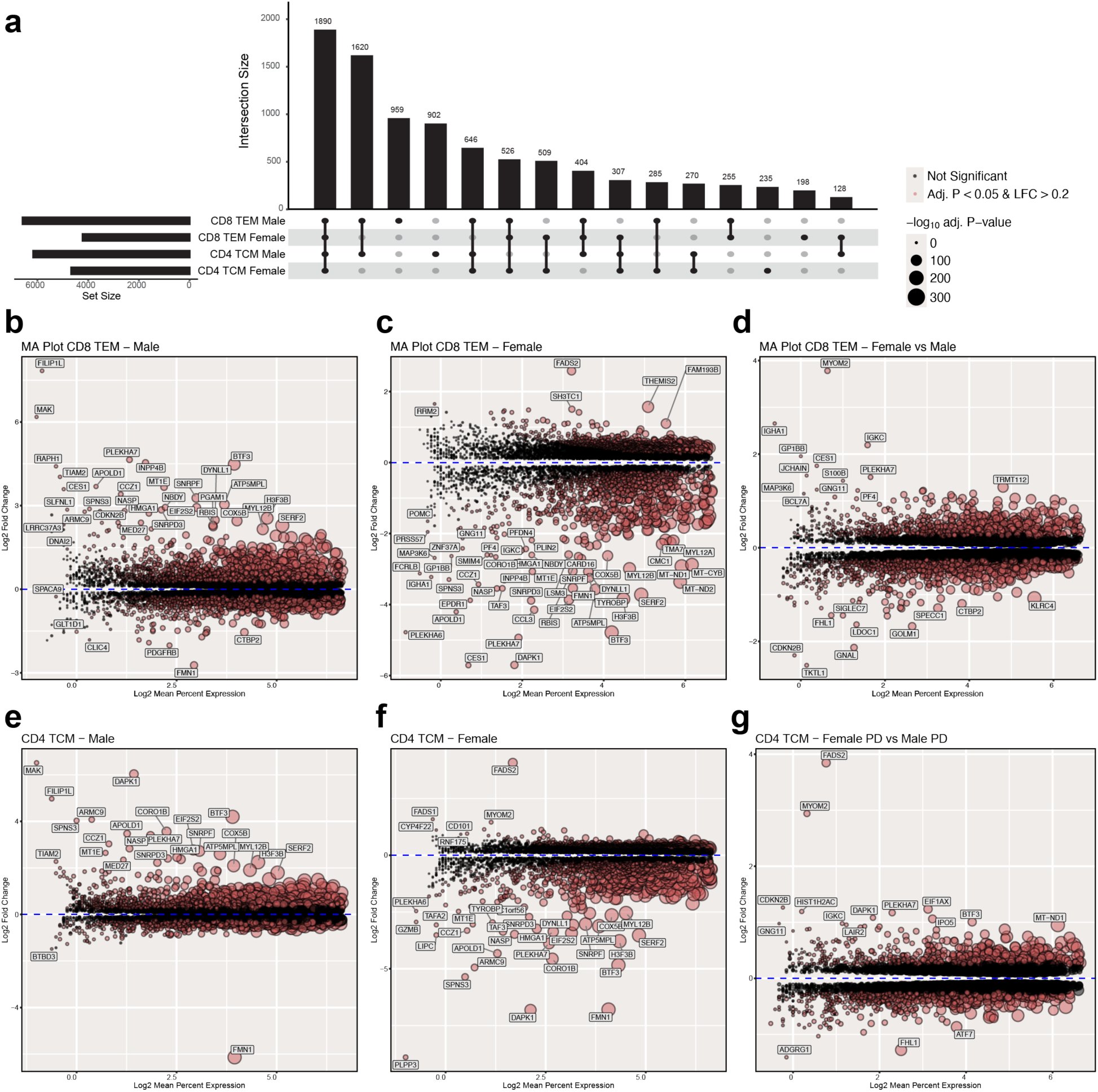
Transcriptomic comparison of differentially expressed genes (DEGs) across sex and disease status in CD8+ effector memory T cell (Tem) and CD4+ central memory T cell (Tcm). **a**) UpSet plot of DEG sets in PD that are unique in 2 cell types in males and females and sets in common have two or more cell sets. b-d) MA plots show DEGs in CD8+ Tem in **b**) males **c**) females and **d**) PD females vs. PD males. Positive Log2 Fold Change are genes enriched in PD males and PD females compared to control males and control females, respectively (**b-c**). Positive Log2 Fold Change indicates genes enriched in PD females and negative Log2 Fold Change indicates genes enriched in male PD. (**d**). **e-g**) MA plots show DEGs in CD4+ Tcm in **e**) males **f**) females and **g**) PD females vs. PD males. Positive Log2 Fold Change indicates genes enriched in PD females and negative Log2 Fold Change indicates genes enriched in male PD. Statistically significant genes are colored red, and the size of the circle corresponds to -log10(Bonferroni-adjusted p-value).

**Figure 8:**
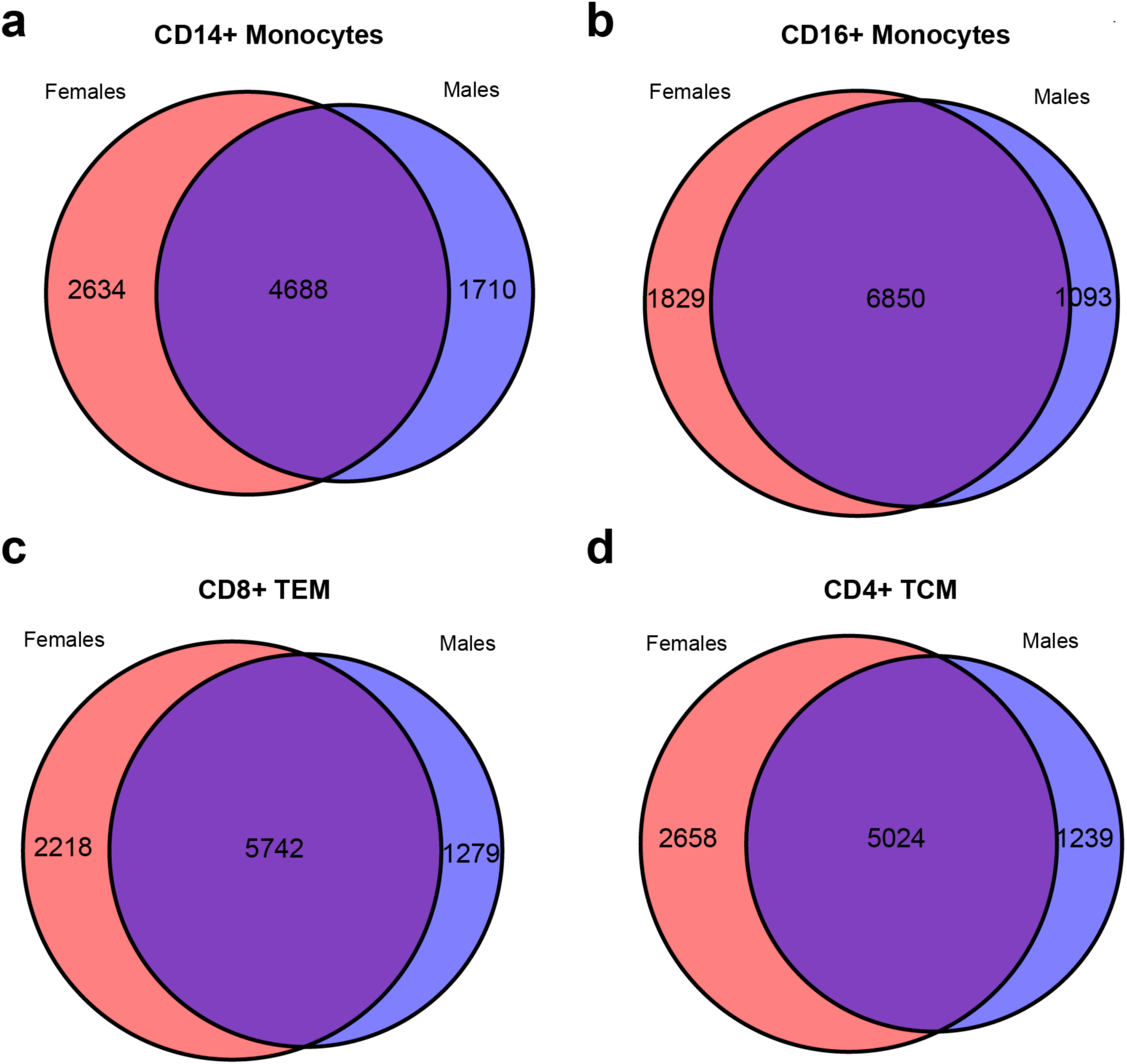
Differential gene expression analysis segmented by sex reveals sex-specific transcriptomic signatures in Parkinson’s disease. Venn diagrams of DEGs that are female-specific (red), male-specific (blue), and shared (overlap) in **a**) CD14+ monocytes **b**) CD16+ monocytes **c**) CD8+ Tem, and **d**) CD4+ Tcm.

There is overlap in DEGs in PD compared to controls in the cell types when parsed by sex, revealing that there are transcriptional shifts unique to PD, but also transcriptional shifts that are unique to PD females or PD males only in these cell types (Fig. 8). Shared and sex-specific DEGs provide an opportunity for understanding sex-related pathology differences in PD.

## Discussion

We leverage high-resolution single-cell transcriptomic strategies to analyze a patient cohort powered to analyze sex as a variable. Our observations are consistent with recent reports indicating significant alterations in immune cell states in PD. Upset plots in Figures 2A, 5A, and 7A summarize both shared lineage-specific changes we identified across the immune landscape. Supplemental tables 3-8 provide a comprehensive list of the specific genes we identified as significantly altered in association with PD across major immune cell types. Overall, our study supports the growing appreciation of the role of the peripheral immune system in PD, while highlighting new targets for foundational and translational studies aimed at identifying novel biomarkers and developing therapeutic strategies for PD.

PD is complex, and patient populations vary widely in their personal habits, treatment strategies, and approaches to the management of clinical progression. This study complements and expands on a similar report published by Xiong et al., which included a scRNA-Seq study of 2 control and 4 PD patients with a total of 48,500 cells analyzed^28^. They report further studies enhanced NK activity in PD patient FAC-sorted cells and tissues that validated their scRNA-Seq findings. This work describes pathway analyses indicating that crucial upregulated NK cell DEGs were associated with immune regulation and cytokine responses. Our work on over 180,000 cells supports the finding of alterations in NK cells and highlights additional pathways of interest, including regulation of Tor signaling and the response to starvation (Fig. 2). Collectively, these convergent transcriptomic findings, framed in a significant body of work *in vitro* and *in vivo*^55–59^, prompt continued evaluation of NK cells in PD.

Gene ontology analysis revealed numerous pathways of interest suggestive of enhanced metabolic and immune response activity consistent with prior findings reported by others. Our study also revealed previously unreported changes. We identified changes in genes related to endoplasmic stress and protein localization of organelles in B cells. ER stress and alterations in organelles, including those of the endolysosomal system, are common features of PD, but research is largely focused on neurons and glia. It is unclear why such changes would occur in B cells; however, it could be speculated to be associated with immunoglobulin production, as this is a primary function of this lineage. ER stress in these B lymphocytes would also align with the evidence for a dysfunctional humoral response during progression of PD, as α-syn autoantibody titers decline as PD progresses^60^. Studies have evaluated the presence of anti-Myelin Associated Glycoprotein (MAG), anti-Myelin Basic Protein (MBP), and anti-synuclein in PD^61,62^, but the origins of these autoantibodies remain unclear. There is evidence of major histocompatibility complex alleles associating with risk of PD^63^, as well as T cell recognition of α-syn peptides in PD^64^. There are greater autoantigenic responses to PINK1 in PD compared to healthy controls, especially in PD males^65^. Analysis of the gene expression profiles in our dataset is anticipated to shed light on this biology with the potential to identify new cellular mechanisms that underlie the observation of autoantibodies.

We previously reported a significant increase in non-steroidal anti-inflammatory drug (NSAID) use among PD patients is associated with changes in immune function, namely, elevated plasma indicators of inflammasome activity^66^. Our prior work was based on data obtained from self-reported surveys in a larger population of PD patients of the same source as this scRNA-seq cohort. Here, we quantified deidentified medical records to annotate medication use among the sequencing cohort. The population reported here is not designed to withstand statistical analysis of medication use; however, Table 2 highlights several noteworthy categories and once again suggests increased use of anti-inflammatory drugs, including NSAID, in PD. As expected, PD patients also more frequently utilized neuromodulatory agents, including dopamine (DA) modulators and Levodopa (alone or in combination with carbidopa; the gold standard of care in pharmaceutical management of Parkinson’s disease), compared with controls. Medication use was comparable between male and female subjects, although fewer males tended to utilize anti-inflammatories and Levodopa compared with females in the PD group. As new high-resolution datasets emerge, it will be important to continue recording and evaluating clinical covariates to ensure that eventual meta-analyses are powered to determine the significance of the potential interactions suggested by the current knowledge base.

Large-scale GWAS studies support deeper analysis of immune cells in PD. The meta-analysis of 17 GWAS datasets (containing PD patients and healthy controls) identified risk loci with probable impact on genes directly associated with immune function. These genes include *NFKB2* and *NOD2*, which are important mediators of the host immune response to DAMPS^26^. *NFKB2* and *NOD2* are expressed in monocytes and function in signaling axes that drive the expression of pro-inflammatory cytokines known to be elevated in PD^67,68^. These include IL6 and IL1B, a canonical target of the inflammasomes^69–72^, a pathway we and others increasingly associate with PD ^73–75^. Figures 4 and 5 indicate significant changes in the monocyte lineage that are supported by prior studies. The aforementioned GWAS analysis identified numerous loci predicted to impact monocyte genes, including *SCARB2*, *VAMP4*, and the familial PD gene *LRRK2*. Other studies point to a change in the monocyte phenotype in PD patients. Differential gene expression analysis in a bulk RNA sequencing study revealed significant alterations in the expression of genes involved in immune activation, such as *HLA-DQB1, MYD88, REL*, and *TNF-α,* suggestive of an alteration in monocytes associated with early-stage PD^76^. A flow cytometry study indicated reduced viability in PD monocytes and altered responsiveness to pro-inflammatory stimuli^77^. A study that negatively selected monocytes from PBMC preparations identified more robust changes in female PD patients^31^. Another study indicated that CCR2^+^ peripheral monocytes were required for inflammation in a viral α-synuclein PD mouse model^16^. These data are concordant with the conclusions of Moquin-Beaudry et al. that classical monocytes in PD are in an activated state, reflected by changes in nucleotide biosynthesis and processing, as well as in their response to viruses, a heightened innate immune state characterized by interferon signaling, cytokine secretion, and antigen presentation machinery^27^. These data also agree with the prevalence of *LRRK2*-associated Parkinson’s disease, as *LRRK2* is known for its role in regulating monocyte activation states, including type I interferon responses, TLR activation, metabolic reprogramming, and antigen presentation^17,78,79^. Our study is significant because high-resolution characterization of the monocyte transcriptome and other immune cells clearly indicates lineage-specific changes in PD, identifying gene-expression programs that may interact with genetic and environmental factors modifying PD risk.

Sex-specific transcriptional profiling of CD14+ monocytes revealed divergent immune and metabolic programs that may influence peripheral inflammatory contributions to PD. Male and female PD monocytes are expressing the PD-associated genes from aforementioned studies differently: *SCARB2, VAMP4, LRRK2, NFKB1, and NFKB2* are each downregulated in male CD14+ monocytes; *SCARB2* and *VAMP4* are not differentially expressed and *NOD2*, *NFKB2*, and *LRRK2* are upregulated in female PD CD14+ monocytes. Female monocytes showed upregulation of genes involved in lipid metabolism, migratory capacity, and immune activation (Fig. 4, 5). These genes indicate a shift towards immune surveillance and regulatory feedback. The concurrent enrichment of lipid metabolic enzymes, pattern-recognition receptors, and chemokine and adhesion molecules suggests female monocytes adopt a metabolically flexible and immunomodulatory activation state that limits excessive inflammatory signaling. Female PD monocytes are activated, but not necessarily canonically inflammatory, conflicting to some degree with the results of Carlisle et al, previous mentioned. Although, both this study and theirs reveal enriched cell migration patterns in female PD monocytes^31^. These slightly divergent results in activation state are possibly related to the covariate differences between the study populations, as the female PD patients in their cohort were within two years of diagnosis, while the patients in this study have an average disease duration of roughly seven years; patients earlier in disease are more likely to be younger and naïve to dopamine-modulator therapies, which may impact monocyte phenotypes as a dopamine-responsive cell type. There is also reported shifts in peripheral immune cell type proportions and states as the disease progresses^47^. These differences highlight the importance of understanding changes in immune phenotypes throughout the progression of PD, and how sex influences these differences. The flexible phenotypic features of female PD monocytes in this cohort could reflect adaptations to reduced estrogen levels after menopause, as estrogen modulates monocyte activation, interferon responsiveness, and metabolism paradigms. Male monocytes, however, exhibited upregulation of genes associated with OXPHOS, translational control, and cellular machinery. These changes reflect a bioenergetically enhanced and stress-primed state, aligning with previous findings^27,80,81^. In males, persistent androgen signaling could reinforce a transcriptionally active state by promoting mechanistic target of rapamycin (mTOR) activation and the downstream inflammatory cascade. Before accounting for sex, PD monocytes are downregulating genes associated with response to virus, a gene ontology pathway commonly linked to interferon stimulated genes (Fig. 4e), and when accounting for sex, DEGs of interferon-stimulated genes appear to be downregulated in males in CD14+ and CD16+ monocyte subsets compared to male controls (Fig 5b, 5e). Compared to females with PD, however, male PD CD14+ and CD16+ monocytes are enriched in these interferon-response genes. Enrichment of OXPHOS and depletion of type I interferon signaling in male PD monocytes replicates the data presented by Carlisle, et al^31^. Together, these data suggest that sex hormones and their metabolic effects shape distinct sex-specific monocyte activation states in PD; females exhibit a more immune-regulatory, balanced phenotype, while males exhibit an oxidatively-driven inflammatory profile. The downregulation of interferon genes in male cells may appear anti-inflammatory, but many of these genes are regulatory, and may leave other inflammatory pathways dysregulated and over-active (for review ^82^). There is also evidence that covariates influence sex-based phenotypes in PD monocytes and these phenotypes may change throughout the course of the disease.

In CD8+ Tem, female cells were characterized by transcriptional signatures of enhanced activation and cytotoxic potential, upregulation of mitochondrial and lipid metabolic genes, and RNA-processing/translation machinery (Fig. 6, 7)^83^. Together, these programs indicate a shift to oxidative phosphorylation, a transcriptionally and translationally active phenotype that favors sustained effector function^84^. Despite activation signatures, these cells are unlikely to exhibit functional cytotoxic capacity. In contrast, male CD8+ Tem cells exhibited more reliance on glycolytic mTOR-associated genes and stress response programs. This suggests a more glycolytic effector program in males tempered by enriched inhibitory signaling, with a loss of typical genes associated with cytotoxic potential. These dimorphic CD8+ Tem states likely have divergent effects on disease biology. Female PD CD8+ Tem programming relies on mitochondrial functioning, vascular interaction, and cytotoxic capacity, and the female CD8+ Tem cells are enriched for T cell activation compared to PD male CD8+ Tem. These traits point to responsiveness to neuroinflammation, neuroinvasion, and possible CNS tissue injury. Male signatures of these cells are associated with heightened metabolic activity, but enrichment of inhibitory genes and depletion of activation-related genes that may dampen overt cytotoxic potential and cell clearance.

These findings synergize with known sex-dependent immune and metabolic patterning across multiple neurodegenerative diseases, highlighting how peripheral immune bias may contribute to divergent clinical trajectories observed between male and female patients. In neurodegenerative diseases such as Alzheimer’s disease and multiple sclerosis, in addition to Parkinson’s disease, females typically exhibit stronger adaptive immune activation, paralleling the OXPHOS-enriched, transcriptionally active CD8+ effector memory phenotype in our female cohort (for review ^85,86^). Enhanced T-cell and microglial activation has been reported in females with Alzheimer’s disease, often accompanied by elevated interferon and cytokine production, which can exacerbate synaptic dysfunction and neuronal loss^29^. In multiple sclerosis, females mount more vigorous T- and B-cell responses, associated with earlier onset and higher relapse frequency, despite slower progression and less total neurodegeneration compared to males later in life^87^. The male PD CD8+ Tem profile in this study highlights glycolytic metabolism and elevated inhibitory receptor expression, mirroring findings in frontotemporal dementia and Alzheimer’s disease where males show greater metabolic stress and neuron vulnerability, but less overt and hyperactive adaptive immune system components^88^. Androgen-driven enhancement of mTOR signaling may play a role in the metabolically demanding state with hyperactive innate cells and stressed adaptive cells^84,89^. These parallels in sex differences to other neurodegenerative diseases suggest a joint contribution of metabolic programming and sex hormone influence in disease.

In Parkinson’s disease, accumulating evidence indicates that the blood-brain barrier (BBB) becomes progressively compromised, allowing increased aberrant peripheral immune infiltration^90,91^, including activated CD8+ T cells and CD14+ monocytes, and subsequent close interaction with the CNS. Infiltrating T cells have been found near degenerating DA neurons in the substantia nigra^92^, and infiltrating monocytes are associated with α-syn-induced neuroinflammation^16^. The female PD CD8+ Tem phenotype may enhance the potential for neurovascular interactions and BBB transmigration, augmenting local inflammation because of their activated state. Estrogen modulates endothelial tight-junction integrity and immune migration^93^, and as it declines post-menopause, may lead to increasing neuro-immune crosstalk. The female CD14+ monocyte phenotype has the potential to contribute further with its enhanced antigen presentation programming. The male CD8+ Tem profile may promote tissue retention and chronic inflammatory signaling within the CNS. Male CD14+ monocytes are primed for pro-inflammatory signaling that is associated with progressive neurodegeneration. These data suggest sex-specific differences in inflammatory signaling, neuroinvasion, and cytotoxic alterations in PD. Sexually dimorphic immune programs may contribute to differential susceptibility, disease trajectory, and neuroinflammatory tone in PD, and present a valuable inroad to understanding differential progression and potential therapeutic targets.

---

**Supplemental Figure 1:**
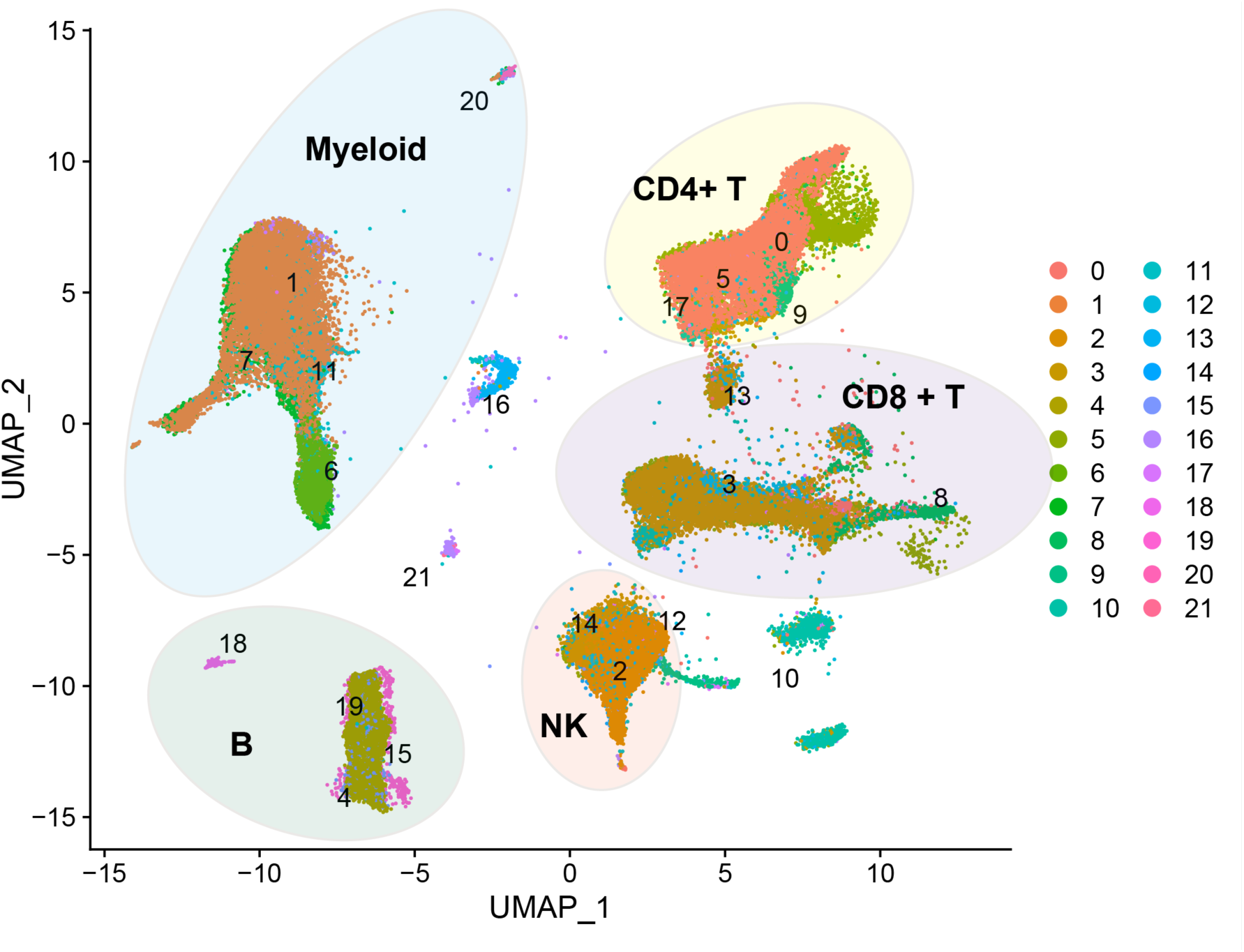
Uniform manifold approximation projection (UMAP) of Louvain clusters. Colored circles represent the major immune cell type each Louvain cluster was identified as (myeloid, CD4+ T, CD8+ T, NK, and B).

